# Diverse deep neural networks all predict human IT well, after training and fitting

**DOI:** 10.1101/2020.05.07.082743

**Authors:** Katherine R. Storrs, Tim C. Kietzmann, Alexander Walther, Johannes Mehrer, Nikolaus Kriegeskorte

## Abstract

Deep neural networks (DNNs) trained on object recognition provide the best current models of high-level visual areas in the brain. What remains unclear is how strongly network design choices, such as architecture, task training, and subsequent fitting to brain data contribute to the observed similarities. Here we compare a diverse set of nine DNN architectures on their ability to explain the representational geometry of 62 isolated object images in human inferior temporal (hIT) cortex, as measured with functional magnetic resonance imaging. We compare untrained networks to their task-trained counterparts, and assess the effect of fitting them to hIT using a cross-validation procedure. To best explain hIT, we fit a weighted combination of the principal components of the features within each layer, and subsequently a weighted combination of layers. We test all models across all stages of training and fitting for their correlation with the hIT representational dissimilarity matrix (RDM) using an independent set of images and subjects. We find that trained models significantly outperform untrained models (accounting for 57% more of the explainable variance), suggesting that features representing natural images are important for explaining hIT. Model fitting further improves the alignment of DNN and hIT representations (by 124%), suggesting that the relative prevalence of different features in hIT does not readily emerge from the particular ImageNet object-recognition task used to train the networks. Finally, all DNN architectures tested achieved equivalent high performance once trained and fitted. Similar ability to explain hIT representations appears to be shared among deep feedforward hierarchies of nonlinear features with spatially restricted receptive fields.

## INTRODUCTION

One of the most striking achievements of the human visual system is our ability to recognise complex objects with extremely high accuracy. Recently, deep neural networks (DNN) using feedforward hierarchies of convolutional features to process images have reached and even surpassed human category-level recognition performance (He et al., 2016; Kietzmann et al., 2018; Lindsay, 2020; Russakovsky et al., 2015; Yamins & DiCarlo, 2016). Despite being developed as computer vision tools, DNNs trained to recognise objects in images are also unsurpassed at predicting how natural images are represented in high-level ventral visual areas of the human and non-human primate brain (Agrawal et al., 2014; Bashivan et al., 2019; Cadieu et al., 2014; Cichy et al., 2016; Devereux et al., 2018; Eickenberg et al., 2017; Güçlü & van Gerven, 2015; Horikawa & Kamitani, 2017; Khaligh-Razavi & Kriegeskorte, 2014; Kubilius et al., 2018; Lindsay, 2020; Ponce et al., 2019; Schrimpf et al., 2018; Xu & Vaziri-Pashkam, 2020; Yamins & DiCarlo, 2016). There is some variability in the accuracy with which different recent DNNs can predict high-level visual representations (Schrimpf et al., 2018; Xu & Vaziri-Pashkam, 2020; Zeman et al., 2020), despite broadly high performance. It remains unclear how strongly network design choices, such as depth, architecture, task training, and subsequent model fitting to neural data may contribute to the observed variations. There are several possible sources that can affect a DNN’s high (or low) correlation with brain representations, and it is important to be able to tease these apart.

First, the architecture of a particular DNN model may cause its representations to be similar to those in the brain. For example, the architecture determines the spatial scale(s) of the image properties able to be represented within each layer. We can gain insight into the importance of such “baked in’’ knowledge by comparing the abilities of different architectures in their random, untrained state (Yamins et al., 2014). In the computer vision literature, deeper architectures have pushed the field towards higher object recognition accuracies (He et al., 2016; Simonyan & Zisserman, 2014; Szegedy et al., 2015), although more recently architectures have been devised that display equal or higher performance with orders of magnitude fewer parameters (Iandola et al., 2016; Sandler et al., 2018). Does depth, across layers or across networks, help predict a model’s correspondence with the brain?

Second, the task training received by a model may have led it to develop computational features that better match those in the visual cortex. It seems intuitive that the success of DNN models at predicting brain data is due in large part to the training the models receive on large datasets of natural images to do behaviourally-relevant tasks such as object recognition (Yamins & DiCarlo, 2016). However, randomly-weighted untrained DNN models are known to explain some variance in visual representations (Cichy et al., 2016; Güçlü & van Gerven, 2015), and at least one study reports higher performance for untrained than trained networks (Truzzi & Cusack, 2020). We can evaluate the contribution of training by comparing the ability of trained and untrained instances of the same architectures to predict brain data.

Finally, two trained models that have learned an identical set of features may nevertheless differ in their apparent similarity to brain representations if they contain different proportions of those features. For example, consider two hypothetical neural network models of low-level visual representations: both networks have learned gabor-like oriented features, but one model contains an approximately equal number of features sensitive to each orientation, while the other has, through a quirk of its training data or task, dedicated almost all of its units to a single orientation. In this idealised example, the representations within both models span the same feature space, but the models will make very different predictions about, for example, how similar the evoked activity in cortical area V1 will be for probe stimuli containing different orientation distributions. We can evaluate to what extent the model has “the right features in the wrong proportions” by measuring how much its predictive power changes after allowing a linear reweighting of its features (Khaligh-Razavi et al., 2017). Many studies reporting high performance of DNNs as models of visual cortex allow linear reweighting of individual features (e.g. Agrawal et al., 2014; Bashivan et al., 2019; Cadieu et al., 2014; Güçlü & van Gerven, 2015; Horikawa & Kamitani, 2017; Ponce et al., 2019; Yamins et al., 2014), while others treat the representations within a layer of a network as fixed (e.g. Khaligh-Razavi & Kriegeskorte, 2014; Truzzi & Cusack, 2020; Zeman et al., 2020). Depending on the particular research question, one may be more interested in model performance with or without fitting to brain data. A good model should have learned features which serve as a good basis set for the features represented by the brain. If one model outperforms another after allowing a linear reweighting of the features within each, then we have reason to think that the features it has learned are a better basis set for the sorts of features computed in the brain.

Here we systematically evaluate the contributions of task training and feature reweighting to the ability of models to predict representations of objects in the ventral stream, across nine state-of-the-art computer vision DNNs. We use representational similarity analysis (RSA) to evaluate the correspondence between fMRI brain activity patterns elicited by viewing object images, and representations of those images in networks. We compare a diverse set of deep neural network models, varying widely in depth (8-201 layers; 25 to 825 processing steps), in terms of their ability to explain the representational geometry in human inferior temporal (hIT) cortex. Each model is tested both in an untrained randomly initialised state and after object-categorisation training. We use principal component reweighting of features within each layer, and reweighting of layers, to best predict the hIT representation. After PCA and hIT-fitting, each model is then tested on its ability to predict the hIT representational dissimilarity matrix (RDM) for an independent set of images in an independent set of subjects. This analysis ensures that the evaluation is not biased by overfitting to either images or subjects.

## METHODS

### Stimuli

Both human participants and neural networks were shown the same set of 62 coloured images depicting faces, objects and places, segmented and presented on a grey background of 427 x 427 pixels (see Figure 1a). The image set was constructed to include a balance of animate (faces and bodies) and inanimate (objects and scenes) stimuli, with animate stimuli further divided into human and animal faces/bodies, and inanimate stimuli divided into man-made and natural objects/scenes. Of the 20 human face images, 14 (7 male, 7 female) were closely matched for low-level image statistics, depicting faces in a 30 degree semi-profile view with matched lighting and matched color-histogram profiles. The remaining face and non-face images contained greater pose and image variation, and were a subset of those previously used in (Kriegeskorte, Mur, Ruff, et al., 2008) and (Kiani et al., 2007).

**Figure 1.**
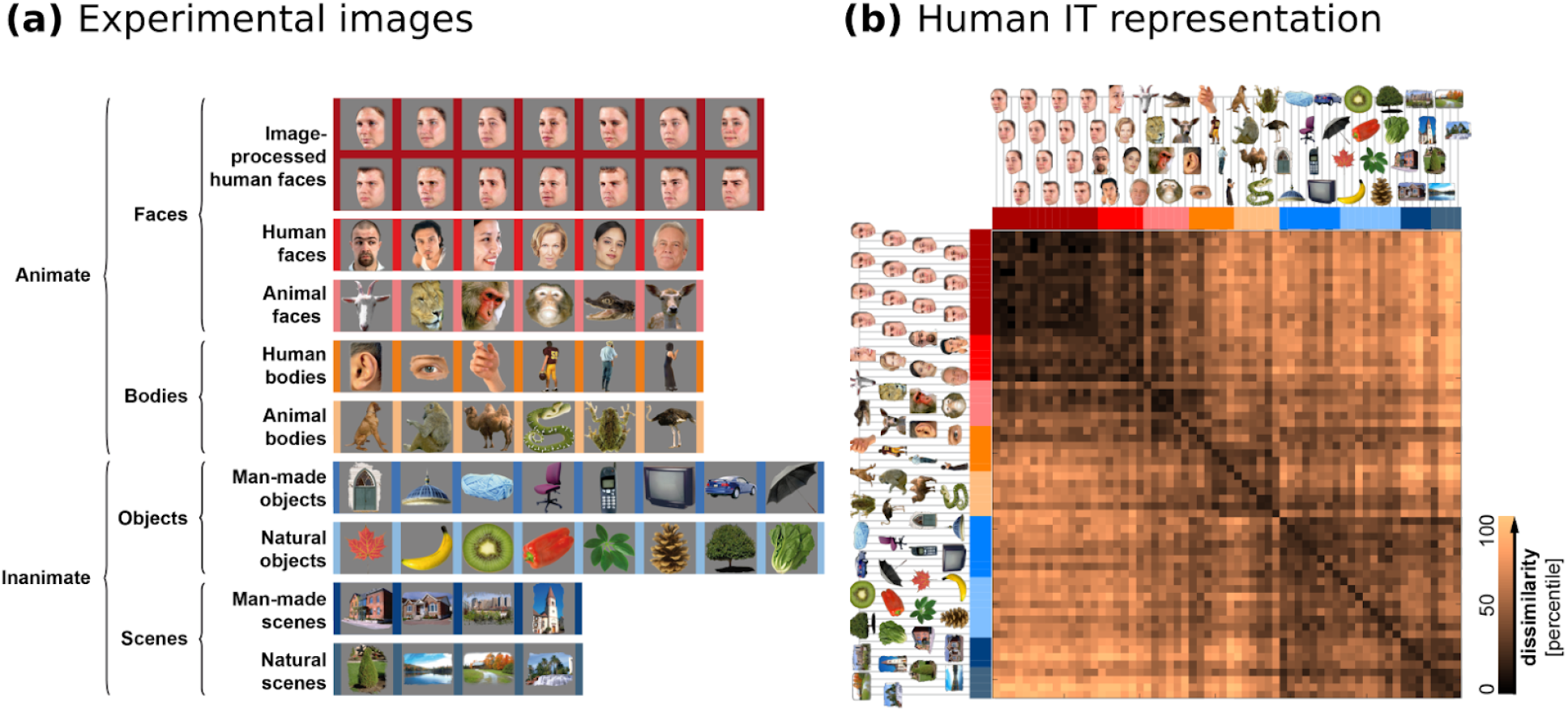
Human fMRI dataset. Sixty-two stimulus images **(a)** were shown to 24 human participants in a rapid event-related fMRI experiment. A representational dissimilarity matrix (RDM) was constructed for each participant from the crossvalidated mahalanobis distances between the multi-voxel activation patterns elicited by each image in inferior temporal cortex (hIT). The average of all individual RDMs is shown in **(b)**, with dissimilarity between the hIT representation of each pair of images expressed as a percentile for visualisation purposes.

### Human fMRI procedures

Deep neural networks were tested against a pre-existing human fMRI dataset (Walther, 2015; Walther, Diedrichsen, et al., 2016), described below.

#### Participants

Participants were 24 healthy adult volunteers (15 female) naïve to the goals of the experiment, with normal or corrected-to-normal vision. Participants gave informed consent and the study was approved by the MRC Cognition and Brain Sciences ethical review panel and conducted in accordance with the Declaration of Helsinki.

#### Image familiarisation

Before the main fMRI experiment, participants were familiarised with the stimuli and task outside the scanner by completing five runs of the one-back task (i.e. five complete cycles through the stimulus set). Participants were also instructed to pay attention to the 62 images shown, and try to commit them to memory. After completing the second of two fMRI sessions, their recall of the images was tested. A total of 128 isolated objects on grey backgrounds was shown to participants, of which 62 were the experimental stimuli. During this recall block, images were shown in a random order, with each image repeated twice, and participants were asked to identify which they had previously seen during the fMRI sessions. On average, recognition accuracy was 92% (standard deviation 0.051%), indicating that participants had attended to and learned the experimental stimuli.

#### fMRI task

During the fMRI scans, participants engaged in a one-back task in which they were instructed to press a button if the present image was a repeat of that shown on the immediately previous trial. While performing the task, participants were asked to keep fixation on a central fixation cross. On each trial, one image was presented in the centre of a grey screen, subtending 7 degrees of visual angle (dva). Images were shown for 500ms with a trial-onset asynchrony of 3 seconds. During each run, 56 of the stimuli were each presented once. The other 6 were presented twice, appearing as repeats in the one-back task. Stimulus order was randomized within each run for each participant. The stimulus sequence also included 30 baseline trials with no image stimulus, comprising 5 blank trials at the beginning of each run, 5 at the end, and 20 randomly interspersed within stimulus trials.

#### MRI measurements

Each participant undertook two scanning sessions on separate days, each consisting of 12 functional runs lasting approximately 5 minutes. Functional images were acquired on a Siemens Trim-Trio 3 Tesla MRI scanner with a 32-channel head coil. For each functional run, we recorded 135 volumes containing 35 slices, each using a 2D echo-planar imaging sequence (TR = 2.18 s, TE = 30 ms, flip angle = 78°, voxel size: 2mm isotropic, interleaved slice acquisition, GRAPPA acceleration factor: 2). We also acquired a high-resolution (1mm isotropic) T1-weighted anatomical image in each session, using a Siemens MPRAGE sequence.

#### Data pre-processing

Image preprocessing was performed using SPM8 (https://www.fil.ion.ucl.ac.uk/spm/). For each participant, images from both sessions were jointly processed, after discarding the first two volumes of each run to prevent T1 saturation effects in the baseline of the regression coefficients. Pre-processing consisted of the following steps, in order: slice-scan-time correction, 3D head motion correction by aligning to the first echo planar image (EPI) of the first run of the first session, re-slicing, and co-registration of the high-resolution anatomical images to the session mean EPI. No smoothing was performed.

#### hIT ROI definition

For each participant and hemisphere, visually responsive voxels were first identified based on an independent functional localizer scan in which responses to 432 images of faces, places, objects, and scrambled objects were contrasted against baseline. An anatomical mask was used to select the subset of those voxels that lay within parahippocampal, entorhinal, fusiform, inferior temporal or lateral occipital regions, using the FreeSurfer package (http://surfer.nmr.mgh.harvard.edu/). Any voxels belonging to early visual cortex (V1, V2 and V3) were excluded, as defined using a cortical surface template projected into each subject’s native volume (Benson et al., 2014).

#### Estimating stimulus response patterns

Response patterns were calculated as in (Walther, Nili, et al., 2016), using multivariate noise normalisation to improve the reliability of dissimilarities subsequently measured between response patterns. Beta response weights were estimated using general linear modeling (GLM) with ordinary least squares. Timecourse data of each run were modeled using 62 stimulus predictors, separately for each subject and session. Six additional one-back predictors were included to model repeated image stimuli in the one-back task. For each run, we included 6 head motion predictors (3D translation and rotation coordinates) and one intercept.

Before fitting, the timecourse data and the design matrix were filtered to remove low-frequency trends. As cross-validated dissimilarity estimates, as used here to derive hIT RDMs (details below), require two independent estimates of stimulus response vectors, two sets of GLMs were fitted for each session and subject (Walther, Nili, et al., 2016). For each of the 12 imaging runs, one GLM was fit on the data of the individual run, while another GLM was jointly fit on the data of the remaining runs, thereby keeping the GLM estimates independent. For the latter GLM, data from the included runs was concatenated for each stimulus (i.e. 11 entries per predictor in the design matrix), which stabilizes the regression weights. Nuisance regressors were modeled separately for each concatenated run. Finally, the 62×P stimulus response beta estimates from each GLM were normalised for multivariate spatial noise by the P×P variance-covariance matrix estimated from the time-course residuals, where P is the number of voxels.

#### Estimating representational geometry

To investigate the representations in hIT, we used Representational Similarity Analysis (RSA; (Kriegeskorte, Mur, & Bandettini, 2008; Nili et al., 2014)). RSA characterises the underlying representations of a given system via representational dissimilarity matrices (RDMs), consisting of dissimilarity estimates for all pairs of stimuli. The set of all pairwise distances describes the geometry of response patterns in high-dimensional activation space (Kriegeskorte & Diedrichsen, 2016) and can be used to compare representations across different systems (here, hIT and DNNs).

For each subject, a hIT RDM was computed by taking the cross-validated Mahalanobis distances between the patterns elicited by each pair of images (Walther, Nili, et al., 2016). This distance measure, also known as ‘crossnobis distance’ does not exhibit an additive bias, unlike non-crossvalidated distance measures, and so preserves a meaningful zero-point when two images elicit identical (but noisy) representations (Walther, Nili, et al., 2016). We calculated leave-one-run-out crossvalidated distances using the two sets of response pattern estimates derived from GLMs fitted to non-overlapping imaging runs. Separate RDMs were derived from the left and right hemispheres, and then averaged to create a single hIT RDM for each participant, of size 62×62 (1891 unique pairwise image dissimilarities).

### Deep neural network models

#### Network architectures and training

We investigated nine deep convolutional neural network (DCNN) architectures representing various states of the art from the computer vision literature over the past eight years (see Table 1). The architectures varied widely in the number of unique processing steps they involved (e.g. convolution, nonlinearity, pooling, batch normalisation, concatenation), from 25 sequential processing steps (Alexnet; (Krizhevsky et al., 2012)), to 825 steps with branching nodes and skipping connections (Inception-Resnet-v2; (Szegedy et al., 2017)). Their sizes varied from 1.24 million parameters (Squeezenet; (Iandola et al., 2016)) to 138 million parameters (VGG-16; (Simonyan & Zisserman, 2014)).

**Table 1.** Details of the nine computer vision DCNNs evaluated. *“Imagenet Top-5 Error”* records the percentage of times the correct object label was not in the network’s top five guesses for the public test set of the Imagenet 1000-way object classification database, used in the Imagenet Large Scale Visual Recognition Competition (ILSVRC). Error rates are as reported for a single model and single crop of each test image for a PyTorch implementation of the models (from: https://pytorch.org/docs/stable/torchvision/models.html), with the exception of InceptionResnet-v2, which is the single-model error rate reported in the original publication. Note that these error rates may differ from the model’s ILSVRC result, because competition results are calculated on a confidential test image set, and because competition submissions often use ensembles of networks and/or data augmentation at test time. The column *“Layers Selected for Evaluation”* briefly describes the criteria we used to select key processing steps within each network, in order to evaluate their match to neural representations in human IT cortex.

| Name | Reference | Imagenet Top-5 Error | Depth (layers) | Number of Parameters (millions) | Layers Selected for Evaluation |
| --- | --- | --- | --- | --- | --- |
| Alexnet | Krizhevsky, Sutskever & Hinton (2012) | 20.9% | 8 | 61.0 | 8 key layers: Outputs of each convolutional or fully connected layer (after ReLU nonlinearity). |
| VGG-16 | Simonyan & Zisserman (2014) | 9.6% | 16 | 138.0 | 16 key layers: Outputs of each convolutional or fully connected layer (after ReLU nonlinearity). |
| Googlenet | Szegedy et al., (2015) | 10.5% | 22 | 7.0 | 13 key layers: Outputs of first three convolutional layers, outputs of each branching “inception” module (after concatenation), and output of final fully connected layer. |
| Resnet-18 | He et al., (2016) | 10.9% | 18 | 11.7 | 10 key layers: Output of first convolutional layer, outputs of each “residual block” (after addition), and output of final fully connected layer. |
| Resnet-50 | He et al., (2016) | 7.1% | 50 | 25.6 | 20 key layers: Output of first convolutional layer, outputs of each “residual block” (after addition), and output of final fully connected layer. |
| Squeezenet | Iandola et al., (2016) | 19.4% | 18 | 1.24 | 11 key layers: Output of first convolutional layer, outputs of each “fire” module (after depth concatenation), and outputs of final convolutional and pooling layers. |
| Densenet-201 | Huang et al., (2017) | 6.4% | 201 | 20.0 | 103 key layers: Output of first convolutional layer, outputs of each densely-connected block (after depth concatenation), and output of final fully connected layer. |
| Inception-Resnet-v2 | Szegedy et al., (2017) | 4.9% | 164 | 55.9 | 50 key layers: Outputs of first five convolutional layers, outputs of each “inception-resnet” module (after addition and ReLU), and output of final fully connected layer. |
| Mobilenet-v2 | Sandler et al., (2018) | 9.7% | 53 | 3.5 | 20 key layers: Output of first convolutional layer, output of each residual or downsizing block (after batch normalisation), and output of final fully connected layer. |

All models were implemented in the Deep Learning Toolbox for MATLAB 2019b, and were pre-trained by their original developers on the Imagenet Large-Scale Visual Recognition Competition (ILSVRC; (Russakovsky et al., 2015)) dataset. The ILSVRC training set consists of 1.2 million labelled images and the networks’ task is to categorise images as belonging to one of 1,000 possible object and animal categories. All networks had near-identical training data and training tasks—slight differences were due to updates to the ILSVRC training set and image categories over the years, and to differences in training strategies adopted by different research groups, such as which methods of data augmentation were used. Object classification error of the networks, as quantified by top-5 error rate on the ILSVRC validation set, ranged from 20.9% error rate (Alexnet; (Krizhevsky et al., 2012)) to 4.9% error rate (Inception-Resnet-v2; (Szegedy et al., 2017)). This measure captures the proportion of test images for which the correct object category was not one of the network’s top five guesses. Human top-5 classification error rate for this dataset is thought to be around 5.1% (Russakovsky et al., 2015).

In addition to analysing the trained networks, we also created untrained randomly-weighted versions of the same architectures, by replacing all weights and biases in each network layer by random numbers drawn from a Gaussian distribution with the same mean and standard deviation as the weights or biases in the trained network at the same layer. Analysis procedures were identical for trained and untrained networks.

#### Image preprocessing

Before being input to neural networks, the 62 stimuli used in the fMRI experiment were resampled to the native input size of each network architecture (either 224×224, 227×227 or 299×299 pixels) and the mean RGB pixel value of each network’s training images was subtracted.

#### Layer activations

For each architecture, we selected a subset of key layers to analyse, generally consisting of the outputs of convolutional or fully connected processing steps, after applying a nonlinearity. For architectures made up of “modules” or “blocks” of processing steps within which parallel branches of processing occurred, or over which skipping connections passed, we took as key layers the outputs of each submodule after all its inputs had been concatenated or added. Table 1 provides further detail on the criteria for key layer selection within each architecture. Network activation patterns in response to the 62 stimulus images were recorded for each of these key layers.

### Comparing human and DCNN image representations

#### Ecologically-driven principal component selection

The number of features in a layer varied by three orders of magnitude across layers and networks, from over 1 million unit activations in the early layers of some architectures, to 1,000 in the output layers. Before reweighting features, we therefore used Principal Components Analysis (PCA) to match the dimensionality of the representations across all layers, and to bring the number of parameters to be fitted into a feasible range. For each layer of each network, a PCA was performed to extract the first 100 orthogonal dimensions explaining the most variance in the responses to a set of independent ecologically-representative images. For this, a set of 3,020 images was drawn from “ecoset” (Mehrer et al., 2017), a large-scale vision dataset that mirrors the most common, most concrete nouns in the english language that describe basic level categories (such as dog, cat, table, etc.). Ecoset thereby represents categories that describe physical things in the world (rather than abstract concepts) which are of significance to humans. The image set used to calculate the PCA had no overlap with the experimental test set, and were natural photographs with backgrounds (whereas test stimuli were isolated object images on grey backgrounds).

Based on each of 100 principal components, 100 component RDMs were computed by taking the Euclidean distance between unit activation patterns elicited by each image pair, after projecting those activation patterns onto each principal component in turn. These 100 component RDMs, extracted for each layer of each model, were then fitted to human IT RDMs by cross-validation over both participants and images (see Figure 2).

**Figure 2.**
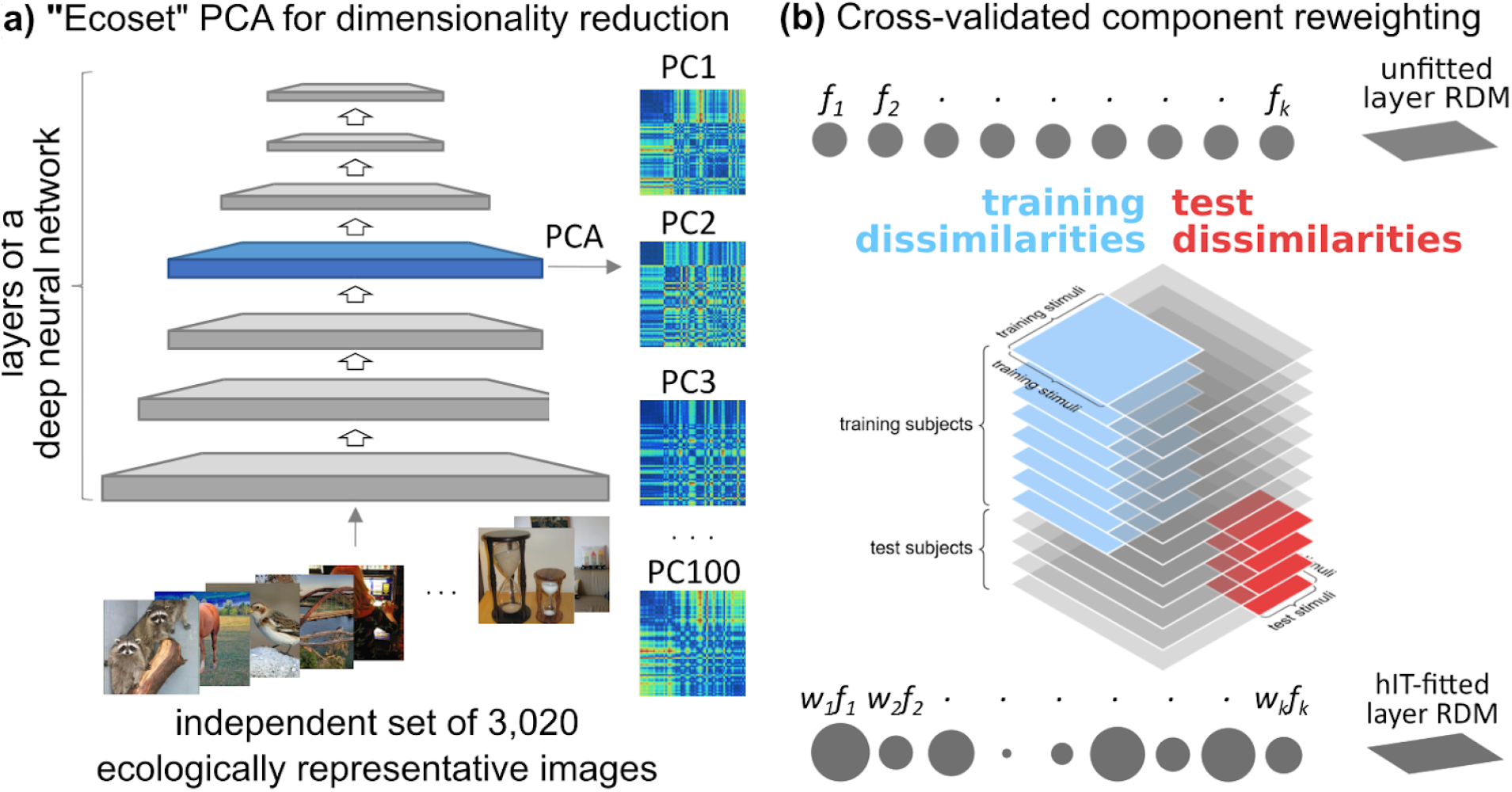
Ecologically-driven dimensionality reduction and component reweighting. An independent dataset **(a)** of 3,020 images derived from object categories that are important to humans was constructed by sampling uniformly from the ecoset dataset (Mehrer et al., 2017). We recorded the activations elicited by these images in each unit of each layer of each network. Based on the data obtained from all units of a given layer, we then ran a Principal Components Analysis (PCA) to identify the first 100 orthogonal components that explained the most variance in the layer’s response to these ecologically more representative images. Based on the projection onto each of the 100 PCs, we then constructed one RDM for the 62 experimental stimulus images. As a distance measure, Euclidean distance between the activation vectors was used. **(b)** After extraction, the 100 component RDMs of each layer were linearly combined within a cross-validated reweighting procedure to predict the human IT fMRI representation (“first-level fitting”). On each cross-validation fold, one non-negative weight was assigned to each component RDM via least-squares fitting. In a second-level reweighting procedure run within the same cross-validation folds, an aggregate prediction for the whole network was then calculated by linearly weighting the per-layer fitted RDMs. All weights were fitted on data from both separate subjects *and* separate image stimuli from those on which they were tested.

#### Cross-validated reweighting

We performed a two-stage reweighting procedure to fit model representations to human IT representations both within and across layers of networks, while cross-validating over both subjects and stimuli (**Figure 2b**). The first-level (within-layer) fitting linearly combined the first 100 principal components within each layer of a model to create a single hIT-fitted RDM for that layer. The second-level fitting then linearly combined these layer-based RDMs across all layers of a network, to create a single whole-network hIT-fitted RDM for that model. The weights for both levels of fitting were estimated within the same crossvalidation procedure. The latter was set up to ensure that model fitting was performed on an independent set of subjects and stimuli. As a result, the reported predictive performance of the DNN models is based on previously unseen subject, and unseen image-data. In addition to model fitting, we calculated the performance of unfitted versions of each layer, and the lower and upper bounds of the noise ceiling, all within the same crossvalidation folds to ensure that all estimates were directly comparable. The full sequence of steps, run for all models within a single bootstrap resampling procedure, is as follows:

1. For each of 1000 bootstrap samples, resample both stimuli and subjects with replacement:

a. For 200 crossvalidation folds:

i. Randomly assign 12 unique stimuli present in this bootstrap sample to be test stimuli. The test set always consists of data from exactly 12 unique images, and typically contains repetitions of some of them. Data from the same image never appears in both training and test sets, even if it is repeatedly present in the bootstrap sample.
ii. Randomly assign 5 individuals present in this bootstrap sample to be test subjects. As with stimuli, the test set consists of data from exactly 5 individuals, and may contain repetitions.
iii. Once training and test stimuli and participants were separated:

1. **Create human target RDM.** Average the data RDMs across training subjects for the training stimuli to create a target RDM to which models will be fit.
2. **First-level (within-layer) hIT-fitting**: For each layer of each model, use that layer’s PCA-derived RDMs as the basis for creating two sets of 100 component RDMs corresponding to the training and test stimuli (these RDMs are subsets of the original RDMs). Use non-negative least squares regression to find the linear weighting of training-stimuli component RDMs that best predicts the human target RDM (derived previously from training participants, only). Create a single-layer predicted RDM for the test images by combining the test-stimuli component RDMs using the weights obtained via the training fit. Compute an unfitted RDM for each layer based on the RDM distances between test images in the original feature space of each layer, with no dimensionality reduction or reweighting applied.
3. **Second-level (across-layer) hIT-fitting**: For each model, take the set of single-layer fitted RDMs calculated at the previous step. Use non-negative least squares regression to linearly weight the first-level fitted RDMs to best predict the human target RDM. As previously, only training RDM distances were used for fitting and the resulting layer weights were applied to the test RDMs to derive a prediction on the test images. This whole-model hIT-fitted predicted RDM aggregates representations across all layers of a network, while allowing the influence of features to be linearly scaled both within and across layers to better match human representations. We also computed a whole-network predicted RDM based on the unfitted per-layer RDMs described in the previous step. This predicted RDM treats each layer as a fixed representation, but allows the influence of each layer on the whole-network predicted RDM to be linearly scaled to better match human data.
4. **Model evaluation**: Evaluate the performance of each of the model predictions as the average Spearman correlation between the predicted RDM and each individual test subject’s RDM for test stimuli. On each cross-validation fold, estimate the performance of the first-level fitted RDMs for each layer of each model, the unfitted per-layer RDMs for each layer of each model, and two second-level (across layer) fitted RDMs for each model, one fitting the first-level fitted RDMs and the other fitting the previously unfitted per-layer predictions.
5. **Noise ceiling calculation**: Calculate the upper and lower bounds of the noise ceiling by taking the correlation between each test subject’s test-stimuli RDM and the mean test-stimuli RDM averaged over either all subjects (upper bound) or the training subjects (lower bound). The noise ceiling provides upper and lower bounds on the expected performance of the true data-generating model, given the inter-subject variability in the data (Nili et al., 2014). By calculating the noise ceiling within the model-reweighting cross-validation folds we ensure that the lower bound estimates are correct for both fitted and unfitted models (Storrs et al., 2020).
b. At the end of the 200 cross-validation folds, average the model performances (first- and second-level) as well as noise ceiling estimates. to create a single estimate of each, for a given bootstrap sample.

## RESULTS

We evaluated how well the representations of object images in each of nine diverse deep neural network architectures could predict those in human inferior temporal (hIT cortex). We analysed performance both for each layer individually and when aggregating across layers in a network. We tested trained and untrained versions of each architecture as well as the effects of allowing linear reweighting of the principal feature components within each layer.

### Object recognition training improves representational correspondence with human IT

First we compared the hIT correlation of every layer in trained and untrained versions of each model (**Figure 3**). We found that training improved representational similarity to hIT, but by a perhaps surprisingly small degree. For each layer and model, we tested whether the distribution of differences in bootstrapped performance between the trained and untrained model contained zero, using a one-tailed test, with an alpha level of 0.05, uncorrected for multiple comparisons. Even using this liberal criterion, only five of the nine models contained layers in which trained performance was better than untrained, and Mobilenet was the only model to show significantly higher performance in most layers after training (see blue asterisks in **Figure 3**).

**Figure 3.**
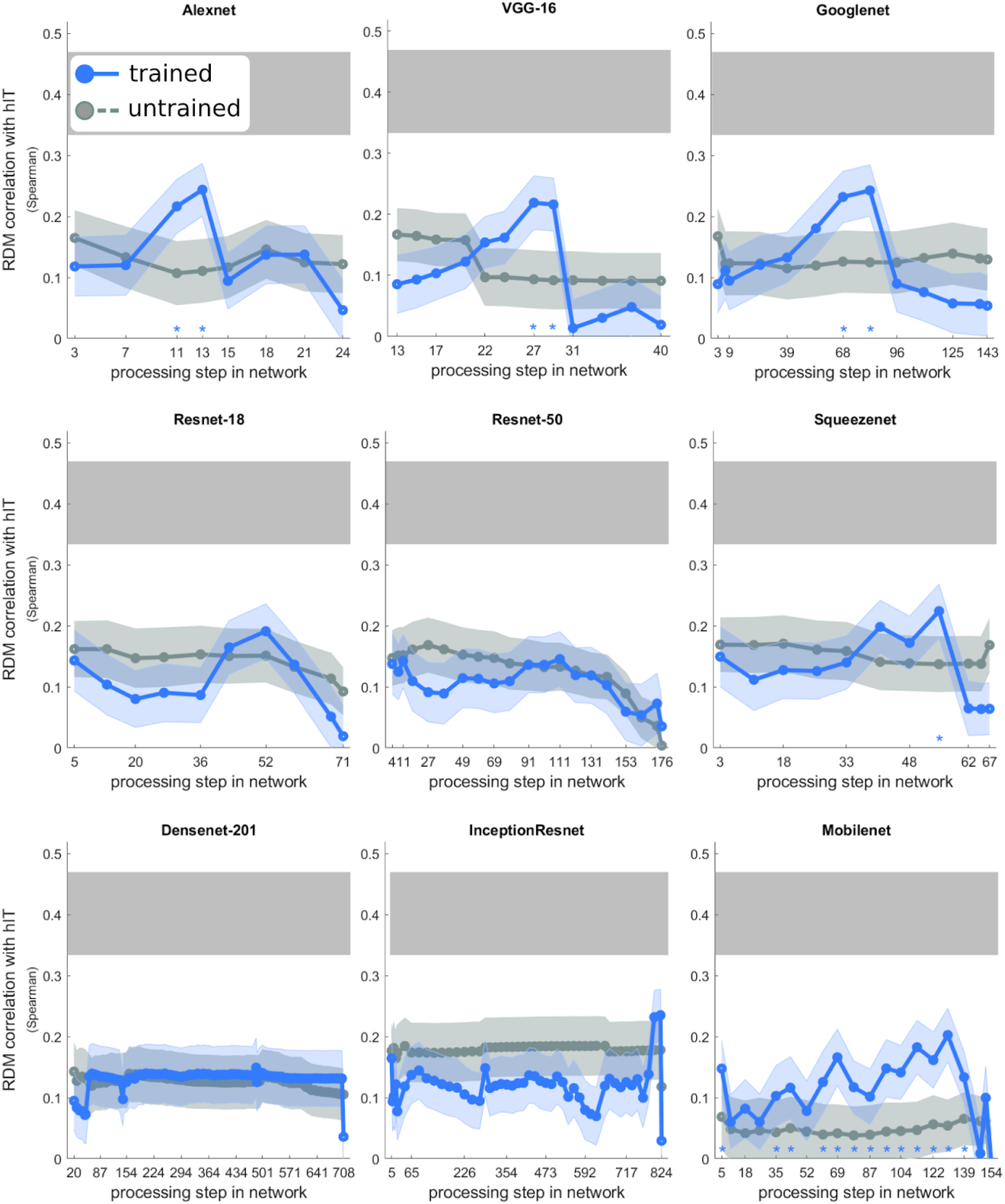
Object recognition training improves hIT correspondence. Panels show the hIT correlation of the full representation in each layer of each architecture, with no dimensionality reduction or feature reweighting, for an object-recognition trained instance of each network (blue) and for a corresponding untrained instance with randomly initialised weights (grey). Each dot corresponds to one of the key layers selected for analysis (see Table 1), and indicates the mean of a distribution of 1,000 bootstrap estimates of cross-validated layer performance, bootstrapped over both subjects and stimuli. For comparability across the diverse architectures, the ‘depth’ of each layer is indicated in terms of the number of unique processing steps up to this point, such as convolution, batch normalisation, pooling, or nonlinearity. Shaded regions indicate the standard deviation of the bootstrap distribution. The horizontal grey bar indicates the lower and upper bounds of the noise ceiling. Blue asterisks above the x-axis indicate that the representation in the trained network performs significantly better than that in the untrained network in this layer (*α* = 0.05, uncorrected). Layers which do not perform significantly below the lower bound of the noise ceiling are indicated by “ns” within the noise ceiling; for all others this comparison is significant.

Notably, however, whereas untrained models showed similar hIT correlation across all their layers, the performance of trained models peaked for processing steps about ½ to ¾ of the way from network input to output. This echoes previous findings of graded similarity between DNN representations and the human ventral stream (Güçlü & van Gerven, 2015; Khaligh-Razavi & Kriegeskorte, 2014; Xu & Vaziri-Pashkam, 2020; Yamins et al., 2014; Zeman et al., 2020), and suggests that the learning of natural image features of the correct complexity is responsible for the superior performance of later layers, rather than inherent architectural properties such as the spatial scale of the representations. Most models show a sharp decline in hIT match towards the final output layers, likely because their training target, a sparse vector indicating which of 1,000 objects possible within the Imagenet ILSVRC challenge dataset is present in an image, forces late layers not to represent graded similarities between images. There was substantial variation among models in how well the best layer correlated with the representation in hIT, but no model explained all of the explainable variance in the human data. A model could be considered to explain all explainable variance in a dataset if it predicts human data as well as individual human subjects can predict the data of other subjects, as quantified by the lower bound of the noise ceiling (Nili et al., 2014; Storrs et al., 2020). For each layer of each model, we tested whether the bootstrap distribution of differences between the lower bound of the noise ceiling and the layer performance contained zero (one-tailed, *α* = 0.05, uncorrected for multiple comparisons). For all layers of all models, the lower bound of the noise ceiling was significantly higher than model performance. Although object-recognition training improves performance, the distribution of visual features learned by task-trained DNNs does not fully mirror those in human IT.

### Reweighting features within trained layers dramatically improves correspondence with hIT

Next we compared the hIT correlation of each layer of the trained networks, both in its full unfitted state, and after model fitting, i.e. reducing dimensionality via PCA on natural images and reweighting the principal components to better predict human representations of held-out data(images and participants, see **Methods**). Reducing and reweighting the feature space improved the correlation of a layer’s representations with that in hIT for virtually all layers of all networks (**Figure 4**). The gradient of similarity between hIT and mid-to-late network layers was enhanced after fitting, which suggests that later layers of task-trained DNNs outperform early layers because they contain features that better match those in late ventral visual cortex in terms of their complexity, structure, or spatial scale, rather than because the preponderance of different features happens to match the brain better than in lower layers. Five of the nine models contained at least one layer in which hIT correlation, after fitting, was not significantly lower than the lower bound of the noise ceiling (based on the bootstrapped distribution of differences, one-tailed, *α* = 0.05, uncorrected. Note that correcting for multiple comparisons would *lower* our threshold for considering a layer statistically indistinguishable from the noise ceiling, and so constitute a more liberal criterion than the test performed here).

**Figure 4.**
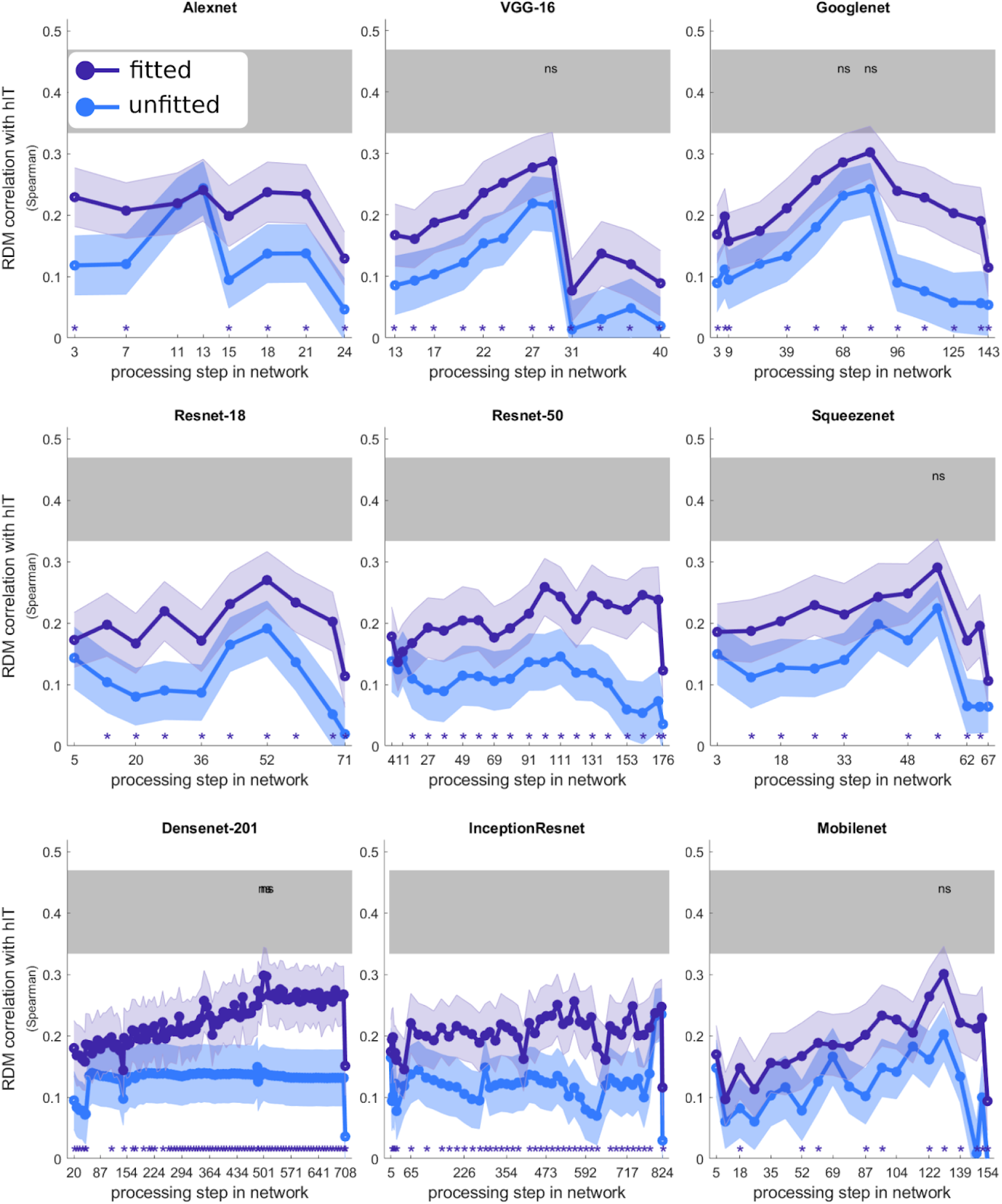
Reducing and reweighting the feature space dramatically improves hIT correlation across most layers in all network architectures. Each panel shows, for one model, the hIT correlation of the unfitted representation in the full original feature space (pale blue lines, same data as shown in Figure 3), and of the same feature space after reducing to 100 dimensions via ecologically-driven dimensionality reduction (see Figure 2) and linearly reweighting those dimensions to fit the hIT representations (crossvalidated over both subjects and images). Blue asterisks indicate that the fitted layer performs significantly better than the original unfitted feature space. Layers in which the fitted representation is not significantly different from the lower bound of the noise ceiling are indicated by “ns”. *α* = 0.05, uncorrected.

### Deeper architectures are not better models of hIT

Despite the wide range of network depths across the nine architectures, spanning 25 to 825 unique processing steps, we found no compelling evidence that deeper networks were better models of hIT than shallower ones. **Figure 5** shows the hIT correlation of each of the fitted layers (i.e. dark blue lines from **Figure 4**) for all models on the same axes for comparability. Although there was a clear improvement in hIT correspondence across early and intermediate layers *within* each model, the peak layer performance was similar across models of different depths. There was no correlation between depth of architecture and whole-network hIT match after combining representations across all layers via second-level fitting, for either trained and within-layer fitted models (*r* = 0.20, *p* = 0.61, *ns*) or trained and within-layer unfitted models (*r* = −0.07, *p* = 0.86, *ns*). Within the range of deep convolutional neural networks capable of high object-recognition accuracy, it does not appear that greater depth leads to more brain-like representations (cf. Kubilius et al., 2018; Schrimpf et al., 2018).

**Figure 5.**
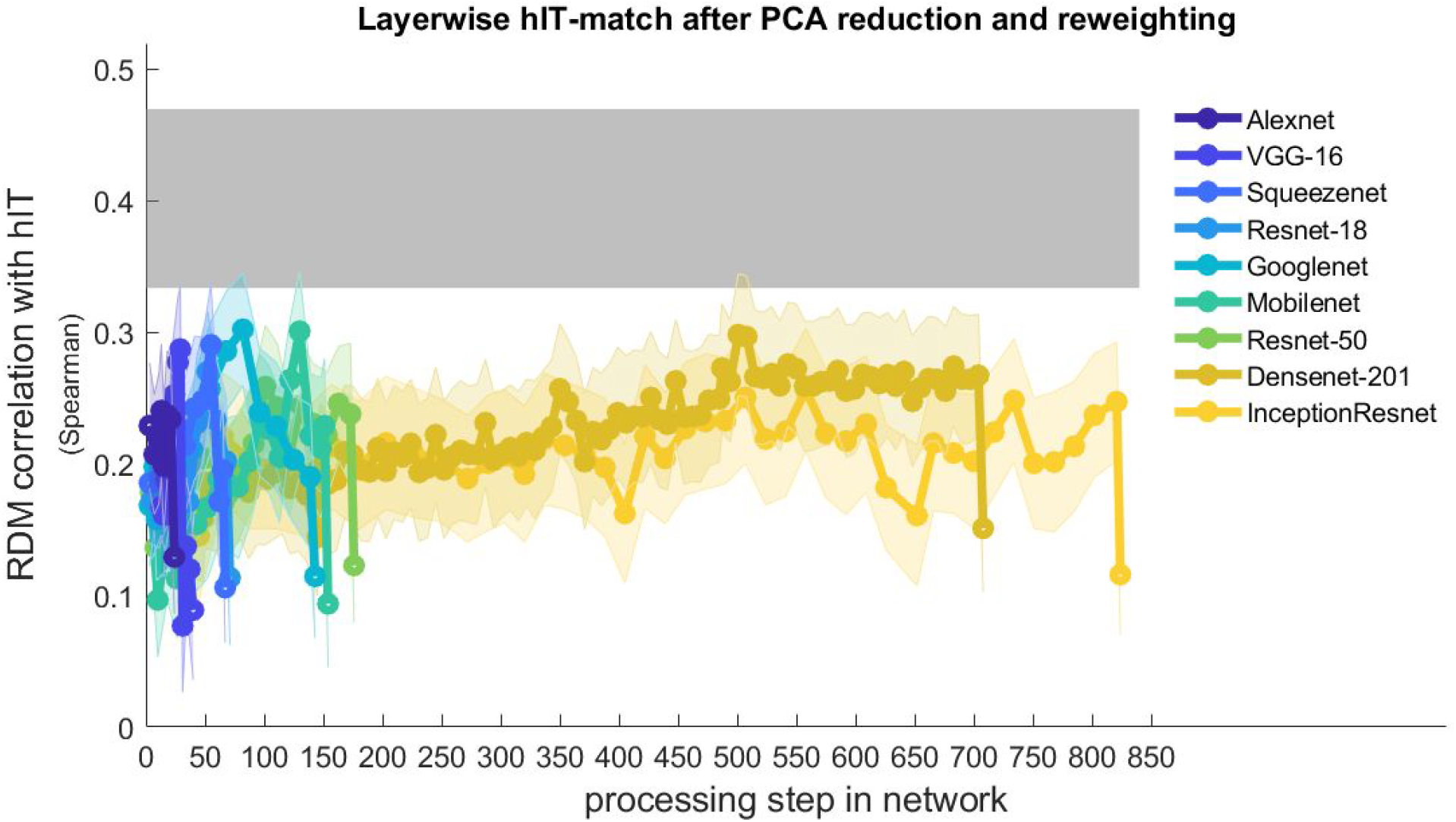
Depth is not the answer. Correlation with hIT representation for all layers of all networks, after “ecologically-driven” PCA reduction and reweighting. Although many models reach representations in their late intermediate layers that well match hIT, increasing depth does not equate to increased hIT match, either within or between architectures, within this range of highly successful object-recognition computer vision models.

### A combination of training and fitting achieves a good match to human IT for diverse deep neural networks

So far we have considered each layer of each network as a separate candidate for predicting hIT. However, it is unlikely that the computational features across large parts of inferior temporal cortex correspond neatly to those in any single processing step of an artificial neural network. We therefore linearly combined the per-layer representational dissimilarity matrices (RDMs) of each network to estimate the hIT correlation of the model considered as a whole, via a second-level fitting procedure (see **Methods**). The inputs to the second-level hIT-fitting were (i) the unfitted per-layer RDMs calculated from the full original feature space in each layer, i.e. without dimensionality reduction and first-level (within-layer) fitting (light grey and blue bars in **Figure 6**), and (ii) the hIT-fitted per-layer RDMs estimated via first-level fitting (comprising both, dimensionality reduction and linear reweighting, dark grey and blue bars). Both levels of fitting were performed within the same cross-validation loops, so that both within-layer and across-layer weights were always fitted based on the same split of training subject and stimulus data and tested on unseen subjects and stimuli.

**Figure 6.**
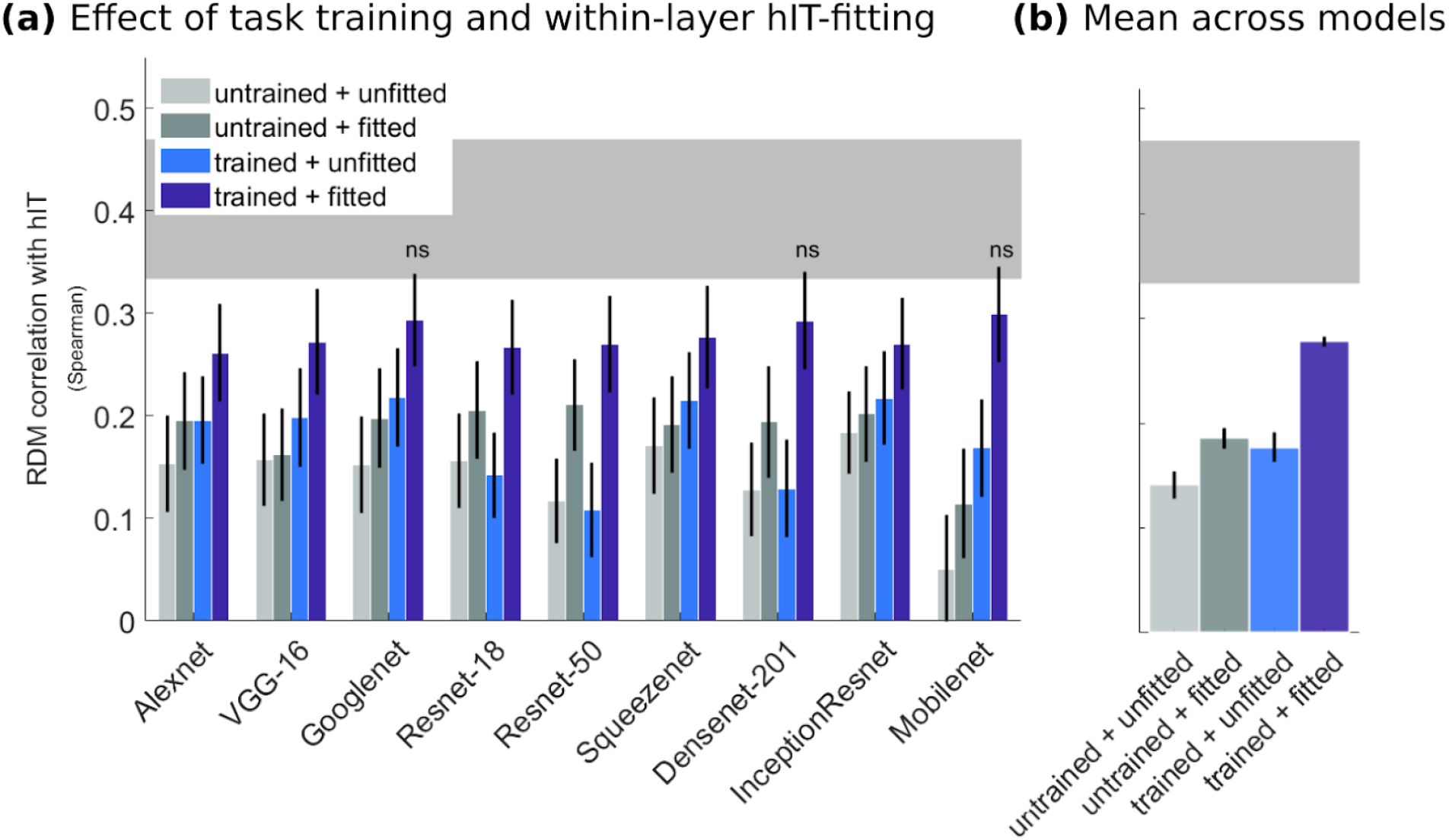
Training and within-layer fitting both improve correlation with hIT. Bars show an estimate of the combined performance of all layers within each of the networks, obtained by second-level (across-layer) fitting in all cases. The fitting procedure is identical to that used to reweight principal components within each layer, except that it takes as input per-layer RDMs rather than per-component RDMs. **Pale grey** bars show the hIT match for the raw (unfitted) feature space of a randomly-weighted instance of the network; **dark grey** bars show hIT match for the same random feature space after PCA reduction and within-layer reweighting; **pale blue** bars show hIT match for the unfitted feature space of the object-recognition-trained network; and **dark blue** bars show hIT match for the trained network after PCA reduction and within-layer reweighting. Error bars indicate the standard deviation of the bootstrap distribution. The horizontal grey bar indicates the lower and upper bounds of the noise ceiling. Models which do not perform significantly below the lower bound of the noise ceiling are indicated by “ns” within the noise ceiling; for all others this comparison is significant, *α* = 0.05, uncorrected. **(b)** Data from (a) averaged across all models. Error bars indicate the standard error of the mean across models.

This whole network analysis revealed that both, object recognition training, and subsequent hIT-fitting improved the correspondence between model representations and those in hIT (**Figure 6b**). A 2×2 (training x fitting) ANOVA treating each of the nine DNN architectures as an independent observation revealed significant main effects of both training (*F*_(1,35)_ = 33.82, *p* < 0.0001), and of fitting (*F*_(1,35)_ = 43.69, *p* < 0.0001). There was also a significant interaction between training and fitting, such that within-layer fitting yields a larger benefit when applied to the features found in layers of trained than untrained models (*F*_(1,35)_ = 6.51, *p* = 0.016).

On average, trained and within-layer fitted models explained 48.0% of the explainable rank variance in hIT representations (squared Spearman correlation 0.08, normalised by the average squared correlations of the upper and lower noise ceilings, 0.16). Starting from an untrained, unfitted model as a baseline, on average across models, task training produced a 57.5% increase in the proportion of explainable rank variance explained (taking the noise-ceiling normalised *r^2^* from 0.12 to 0.19). Similarly, hIT-fitting of the untrained model produced a 73.4% increase (normalised *r^2^* from 0.12 to 0.21). While these are substantial gains, the combination of training and hIT-fitting achieved a superadditive boost. Compared to trained but unfitted networks, reweighting the features within each layer of trained networks led to a further 124% increase in the proportion of explainable rank variance explained (normalised *r^2^* from 0.21 to 0.48). The superadditive interaction between training and fitting suggests that training on the Imagenet object-recognition task causes models to develop features that capture aspects of real-world images that are important to their representation in hIT, but does not cause models to learn the relative prevalence of these features seen in brain data.

After training networks to classify objects, and linearly reweighting their learned features within and across layers, all nine DNN architectures yielded good models of hIT, explaining most of the explainable variance in the data (dark blue bars in **Figure 6a**). For three architectures (Googlenet, Densenet and Mobilenet), the trained and hIT-fitted model was not statistically distinguishable from the lower bound of the noise ceiling, indicating performance on par with the ability of individual human brains to predict representational dissimilarities in other human brains (one-tailed test of whether the bootstrap distribution of differences contained zero, *α* = 0.05, uncorrected). In order to evaluate the relative contributions of architectural differences to model performances before and after reweighting, we ran a permutation test comparing all pairwise differences among trained but only second-level fitted (i.e. within-layer-unfitted) models to those among fully fitted models. Differences among the hIT correspondence of unfitted models were larger than those among the same models after they had undergone within-layer principal components reweighting (Cohen’s *d* = 1.49, *p* = 0.001).

In order to gauge whether the remaining differences in performance among trained and fitted models were likely to be of theoretical interest, we performed an equivalence test (Lakens, 2017) in which we defined the variance in estimates of the lower bound of the noise ceiling as a naturally occurring variation in predicting individual human data. If a difference between two models falls outside the 95% confidence interval of this distribution, the models are more different from one another than human subjects are from one another, which could be considered a threshold for the minimal potentially interesting model difference. For each pair of models, we tested whether the observed differences between the hIT correlation of the two models, across bootstrap samples, were significantly larger than the bound of the 95% confidence interval on noise ceiling variation, and at the same time significantly lower than the upper bound of the confidence interval (Lakens, 2017). After training and hIT-fitting, the differences between models proved statistically equivalent to the differences between human subjects for all models (*α* = 0.05, Bonferroni corrected for 36 pairwise comparisons among models). However, there were larger differences among pairs of trained but unfitted models, with differences in 8 of the 36 pairwise comparisons falling significantly outside of the variability in the lower noise ceiling.

### Finding dimensions of natural-image variation within layers improves correlation with human IT

Our first-level (within-layer) fitting procedure consists of two steps, both of which potentially change the representational geometry: first, the full feature space of a layer is reduced to the first 100 principal components accounting for the most variation in that layer’s activation patterns to natural images of ecological validity, and second, those principal components are linearly reweighted to best predict dissimilarities between images in human IT. In order to assess the relative contribution of each step towards the improvements gained by fitting, we calculated the performance of a version of each model that had undergone the first (dimensionality reduction) step, but not the second (hIT-fitting) step. Within the main cross-validation procedure (see **Methods**) we created an RDM for each layer of each network by uniformly combining the layer’s 100 principal component RDMs with equal weights, and used these as inputs to the second-level (across-layer) fitting procedure. The resulting whole-network hIT correlations are shown in the central column of **Figure 7a**, with those shown previously, based on either the full unfitted per-layer representations, or the dimensionality-reduced and hIT-fitted per-layer representations to the left and right, respectively, for comparison.

**Figure 7.**
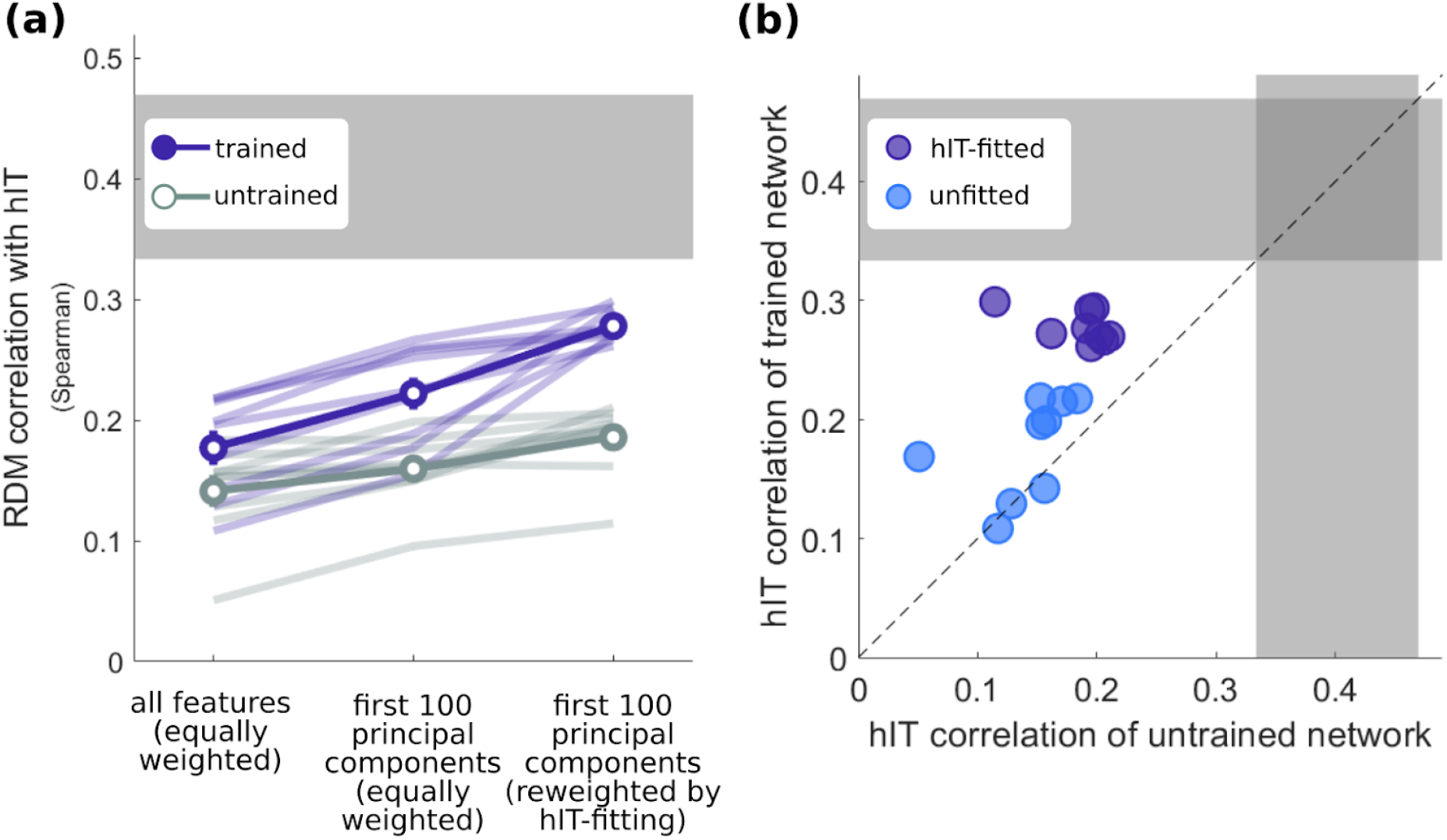
Finding dimensions of natural-image variation within each layer’s feature space improves hIT match. **(a)** Estimates of the whole-network hIT correlation for trained (blue) and random (grey) networks, derived by reweighting layer RDMs obtained from either (left) the full original feature space, (middle) the first 100 components of that feature space, derived via PCA on an independent natural image set, and (right) after reweighting the principal components within each layer to predict hIT representations. **(b)** Estimates of the whole-network hIT correlation for untrained networks (x-axis) compared to their task-trained counterparts (y-axis), derived by reweighting layer RDMs obtained from either (pale blue dots) the full original feature space or (dark blue dots) within-layer hIT-fitted representations.

Despite reducing the dimensionality of the feature space by up to four orders of magnitude for some layers of some networks, from over 1 million units to only 100 principal dimensions, PCA improved the hIT correspondence of models (**Figure 7a**). A 2×2 (dimensionality reduction x training) ANOVA revealed a main effect of both dimensionality reduction (*F*_(1,35)_ = 6.39, *p* = 0.0166) and training (*F*_(1,35)_ = 15.14, *p* = 0.0005). Post-hoc tests on the effect of dimensionality reduction show that models based on the first 100 principal components within each layer matched hIT better than models based on the full feature space, both for trained networks (paired-samples t-test *t*_8_ = 6.87, *p* = 0.0001) and untrained networks (*t*_8_ = 4.84, *p* = 0.0013). There was no interaction between training and dimensionality reduction (*F*_(1,35)_ = 1.09, *ns*). On average, for the trained networks, correlation with hIT could be improved by 25.5% simply by taking dimensions within each layer that account for the most variation in the network’s activations to natural images sampled from a set of object categories that are of significance to the human (visual) experience (Mehrer et al., 2017).

### Differences in hIT correspondence between trained models do not well predict their performance on other model evaluation metrics

After task training, but without per-layer hIT-fitting, there were non-trivial differences among some DNN architectures in how well they predicted representations in human IT (as supported by an equivalence test against the variability in the lower bound of the noise ceiling, described above). Did these differences point to the superiority of certain DNN architectures as models of the brain over others? We investigated whether higher hIT correspondence for unfitted models was associated with higher performance on other model evaluation metrics, such as their performance after within-layer hIT fitting, their ability to predict other neural and behavioural datasets, or their object classification accuracy.

We found that, for within-layer-unfitted models, there was a positive correlation between hIT correlation before and after task training (**Figure 7b**; Pearson Skipped *r*= 0.49, 95% CI = [0.16, 0.96], *α* = 0.05, using the robust correlation toolbox for Matlab (Pernet et al., 2013)). There was no relationship between the performance of trained and untrained models after fitting the within-layer representations of each to hIT (**Figure 7b**; Pearson Skipped *r* = 0.07, *ns*).

Among relatively shallow neural networks, models with higher object classification accuracy tend to provide feature spaces that can better predict the firing rates of neurons in macaque IT to object images (Yamins et al., 2014). Among deeper, higher-performing networks, this effect appears to saturate, and further improvements to classification accuracy no longer translate into higher performance as brain models (Schrimpf et al., 2018). We found no significant association, either among unfitted or fitted models, between accuracy on the ILSVRC object classification task, and correlation with human IT (**Figure 8a**; Pearson Skipped *r* = −0.38 (unfitted), 0.32 (fitted) both *ns*).

**Figure 8.**
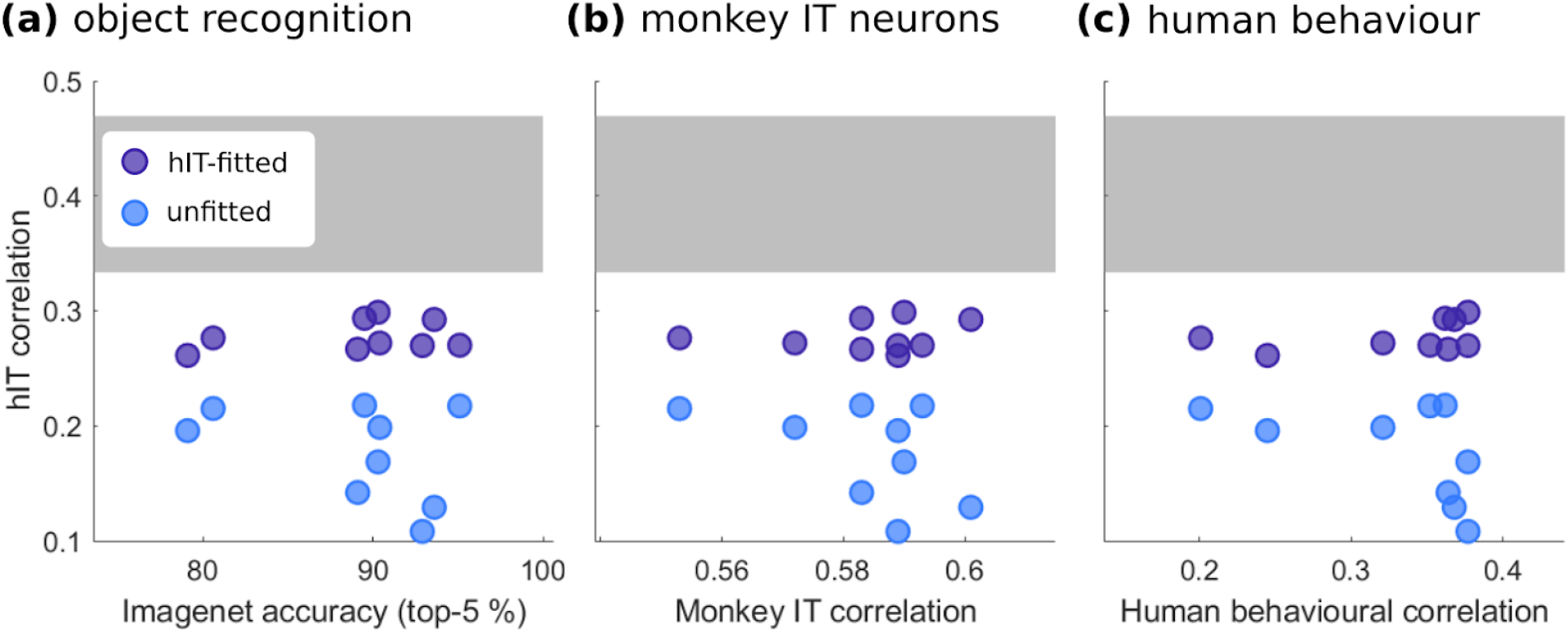
Better hIT models are not consistently better on other model evaluation metrics. **(a)** Relationship between hIT match and object classification accuracy on the ILSVRC dataset for each model (Russakovsky et al., 2015) **(b)** ability of each model to predict neural responses in macaque IT (Majaj et al., 2015; Schrimpf et al., 2018). **(c)** ability of each model to predict categorisation responses in a large set of human behavioural judgements on object images (Rajalingham et al., 2018; Schrimpf et al., 2018).

One of the central goals of computational visual neuroscience is a model that can predict neural representations in visual cortex at multiple levels of granularity, from single neuron responses to the aggregated population signals measured via fMRI, and can also predict the perceptual properties of our visual systems, as measured in behavioural experiments (Funke et al., 2020; Hebart et al., 2020; Rajalingham et al., 2018; Schrimpf et al., 2018; Storrs & Kriegeskorte, 2020). We were therefore interested in whether models that predicted fMRI-based representational dissimilarities in human IT better also predicted electrophysiological or behavioural object perception data better. We compared the hIT correlation of each of the trained models to the previously reported ability of each architecture to predict macaque IT firing rates in response to object images (Majaj et al., 2015; using values reported in Schrimpf et al., 2018), and to predict a set of human behavioural object classification judgements (Rajalingham et al., 2018; using values reported in Schrimpf et al., 2018). Robust correlation analysis revealed no significant relationships with either measure, for either fitted or per-layer unfitted models (for macaque IT **Figure 8b**: Pearson Skipped *r* = −0.47 (unfitted), 0.19 (fitted) both *ns*; for human behavioural judgements **Figure 8c**: Pearson Skipped *r* = −0.52 (unfitted), 0.57 (fitted) both *ns*).

## DISCUSSION

In this work, we investigated a diverse set of DNNs for their ability to predict representations estimated from fMRI data of human inferior temporal cortex. Comparing the predictive performance of untrained and object-recognition-trained network variants we show that task training substantially improves correspondence with representations in hIT. This suggests that training on natural stimuli to perform ecologically relevant tasks yields a set of computational features that form a better basis for predicting those in the human brain. The effect of training is significantly amplified by model fitting, indicating that the relative prevalence of different features in hIT does not automatically emerge from the particular ImageNet recognition task training used to train the networks. Following task-training and two-stage model fitting, the predictive performance of all networks, irrespective of depth, on data from unseen stimuli and participants, was similarly high, explaining 48% of the rank variance in human IT representations. This is similar to the proportion of variance DNNs have previously been found to explain in macaque electrophysiological data (Bashivan et al., 2019; Cadieu et al., 2014; Yamins et al., 2014). The indistinguishable performance levels of diverse architectures, after training and fitting, suggests that the models’ architectural specifics—depth, numbers of feature maps, sizes of filters, presence of skipping or branching connections—matter less than their shared attributes, as relatively deep feedforward hierarchies of convolutional features.

### Training DNNs helps inject domain knowledge, and helps to address why brain representations contain particular features

A number of studies has shown that performance-optimised DNNs can explain representations in high-level regions of human and nonhuman primate ventral visual cortex (Agrawal et al., 2014; Cichy et al., 2016; Güçlü & van Gerven, 2015; Khaligh-Razavi & Kriegeskorte, 2014; Kriegeskorte, 2015; Kubilius et al., 2018; Schrimpf et al., 2018; Xu & Vaziri-Pashkam, 2020; Yamins et al., 2014; Yamins & DiCarlo, 2016). Our results here replicate and extend upon this widely appreciated finding by testing a large variety of models on the same hIT dataset, while at the same time providing estimates for the relative effects of task training and model fitting.

Knowing the size of the effect of task training on a given network’s predictive power serves an important role in our goal of understanding ventral stream computations. Following the normative approach, training models on an external task can help answer the question of *why* the ventral stream computes what it computes (Kietzmann et al., 2018). While a larger set of supervised and unsupervised training objectives needs to be investigated, the present results indicate that exposure to natural images is important for hIT-like representations, as randomly initialised untrained deep architectures performed significantly worse. At the same time, the dramatic benefits of model fitting suggest that training on the ImageNet ILSVRC challenge does not lead to the correct relative feature distribution.

In addition to answering the *why* question, task-trained models are advantageous because they can harness large training datasets that go beyond what can be measured within a given neuroscientific experiment. Vision, like other feats of intelligence, requires knowledge about the world. In particular, recognition requires knowledge of what things look like. In order to explain task performance and high-level responses, therefore, a model needs the parametric capacity to store the requisite knowledge. Direct training with millions of natural images provides a highly efficient way of arriving at useful visual features deep within network hierarchies. Yet, the differences in predictive performance between random and trained versions of the same architecture are surprisingly small, suggesting we need to be cautious about interpreting the features learned by trained networks as being informative about the particular features represented in the ventral stream.

### Reweighting features improves hIT correspondence, and reveals common performance across diverse models

After finding dimensions of important variance, and reweighting to adjust the relative strength of those feature dimensions, good prediction of human IT representations in our dataset could be achieved by all models, with minimal differences in performance among very diverse architectures ranging from 8 to 201 layers in depth. The fitted and unfitted performance estimates provide us with different information, and experimenters may wish to use or not use fitting depending on their modelling objective.

The substantial benefit of hIT-fitting (124% improvement, for trained models) should not be surprising, for at least two reasons. First, the object-recognition-trained networks had as their only requirement the classification of 1,000 nameable objects, with a distribution highly unlike that found in the human visual diet — for example, the ILSVRC categories do not contain “person” or “face” (Mehrer et al., 2017; Russakovsky et al., 2015). The human ventral stream, in contrast, must subserve a wide range of behaviours beyond recognition such as navigation, interaction, and memory. For some research questions, we may be most interested in the performance of models *without* allowing feature reweighting. Model fitting (including encoding models, and single- or two-stage RDM reweighting) always deviates models away from their “native” feature coding, by allowing the prevalence of different features to be adjusted (Khaligh-Razavi et al., 2017). If we are interested in which architectures, training objectives and visual diets give rise to *distributions* of features similar to those found in the human visual system, the unfitted performance of models will be most informative.

A second consideration is that, even if an ideal DNN model of human IT were to exist, containing exactly the features and distribution of those features found in the brain, the measurement processes giving rise to data would bias the prevalence of *measured* features (Kriegeskorte & Diedrichsen, 2016). This provides one motivation *for* reweighting features — since we know that our measurement processes can introduce bias in feature sampling, requiring a model to match the measured prevalence of features might be too strict a criterion. Instead of reweighting existing features that emerge via task-training, researchers have recently started using data from the human ventral stream to directly learn the network features themselves in end-to-end training on natural stimuli (Kietzmann et al., 2019; Seeliger et al., 2019). Such procedures serve the important function of verifying that a given network architecture chosen is in principle capable of mirroring the right representational transitions observed in the brain.

### The future of DNNs as models in visual neuroscience

The advent of high-performing object-recognition DNNs in computer vision has provided visual neuroscience with unprecedentedly good models for predicting visual responses in the human and non-human primate brain (Kietzmann et al., 2018; Kriegeskorte, 2015; Lindsay, 2020; Schrimpf et al., 2018; Yamins et al., 2014). The deep learning modelling framework promises much more, beyond evaluating the effectiveness of off-the-shelf feedforward DNNs trained on computer vision tasks. Despite the achievements of such models, they are not yet perfect models of biological vision, exhibiting fragility in the face of noise and other perturbations (Geirhos et al., 2017, 2020), an over-reliance on textural information (Geirhos et al., 2018), and limited ability to predict brain responses to artificial stimuli (Xu & Vaziri-Pashkam, 2020).

Going forward, deep learning models in visual neuroscience will more broadly explore the space of objective functions, learning rules, architectures (Richards et al., 2019), and training diets (Mehrer et al., 2017). Recurrent networks can recycle neural resources to flexibly trade speed for accuracy in visual recognition and show great promise as models of temporal dynamics in visual cortex (Güçlü & van Gerven, 2017; Kietzmann et al., 2019; Nayebi et al., 2018; Spoerer et al., 2017; van Bergen & Kriegeskorte, 2020). Unsupervised learning objectives provide rich and ecologically feasible ways of getting complex knowledge about the visual world into the brain (Fleming & Storrs, 2019; Storrs & Fleming, 2020). Models should be able to predict both internal representations and behavioural data (Funke et al., 2020; Jozwik et al., 2017; Storrs & Kriegeskorte, 2020), and will be tested using larger datasets, with higher noise ceilings, and with stimuli designed to tease apart the differences between model predictions (Golan et al., 2019). We have come a long way, but are only just beginning to explore the full potential of deep learning in visual neuroscience.

## ACKNOWLEDGEMENTS

This work was supported by the UK Medical Research Council (program MC-A060-5PR20), a European Research Council Starting Grant (261352) to NK, a Vice Chancellor’s Award by the Cambridge Trust to JM, and an Alexander von Humboldt Foundation fellowship awarded to KRS.

